# The Efficacy of the Antimicrobial Peptide Melittin and the Enzyme DNase I Against *E. coli* Biofilms

**DOI:** 10.1101/523159

**Authors:** Abigail Phillips

## Abstract

Biofilms are aggregations of bacteria that often cause nosocomial infections and are tolerant to antibiotic treatments due to a variety of factors, including the physical barrier of their matrix. One field of treatment that has been explored is the use of antimicrobial peptides, like melittin. Melittin has been demonstrated to have antimicrobial effects on bacteria in both a biofilm and a planktonic state. Its required concentration to affect biofilms, however, is too high to be applicable in clinical settings. One potential way to increase efficacy is to use a compound like DNase I that degrades the matrix and thereby increases the penetration of melittin into the biofilm. I investigated whether DNase I would enhance the efficacy of melittin in relatively low concentrations that could be clinically viable. Melittin and DNase I were applied to *E. coli* biofilms in 24-well plates, both individually and in combination; biofilm density was measured at two time-points afterwards, using a crystal violet staining assay. The results showed DNase I was the most effective treatment. Surprisingly, none of the melittin treatments were statistically significant. A second experiment investigated variation among 11 bioreplicates and found that there was approximately 5-fold variation in response to treatment with DNase I. Given these promising results, further study on DNase I should be conducted.

**Importance:** This work is important as it is crucial to have as many viable treatment options as possible for biofilm infections. Biofilm infections are common and many of our standard antibiotic options are simply not effective against them. According to the NIH, 65% of microbial infections and 80% of chronic infections are associated with biofilms, costing the United States millions of dollars each year (Jamal et al., 2018). Every avenue that could potentially further efforts to increase the efficacy and practicality of treatment options should be explored. Antimicrobial peptides are one of the most promising fields for biofilm treatment, and this study further investigates the potential for treatment that they hold.

## Introduction

Biofilms are cooperative groups of bacteria that are highly antibiotic resistant and pose a serious health risk to those who develop infections that include them. They can form rapidly and are extremely difficult to kill or remove from a surface (Figure 1) (Taylor et al., 2014). Specifically, they are composed of one or more species of bacteria clustered together on a surface surrounded by an extracellular matrix. This generally occurs in an environment that is moist and includes a surface for the biofilm to attach to such as many medical instruments, including catheters and heart valves (Jolivet-Gougeon & Bonnaure-Mallet, 2014). Out of 721,800 nosocomial infections in the United States in 2011, 250,800 of those were urinary tract or surgical site infections, both of which commonly include biofilms (Magill et al., 2014). Because biofilms are a form of stress response, their formation is rapid, and once they exist they are very hard to get rid of due to their high resistance to antibiotics and the host immune system (de la Fuente-Núñez et al., 2013). This resistance is a product of a variety of factors including the presence of persister cells, the biofilm matrix, and the general differential gene regulation within the biofilm. One possible way to kill biofilm infections is the use of antimicrobial peptides. These are small molecules that can eliminate a wide variety of biofilms, usually by creating pores in the membranes of the bacteria, and their doses can be clinically achievable, especially in combination with regular antibiotics (Chung & Khanum, 2017).

**Figure 1:**
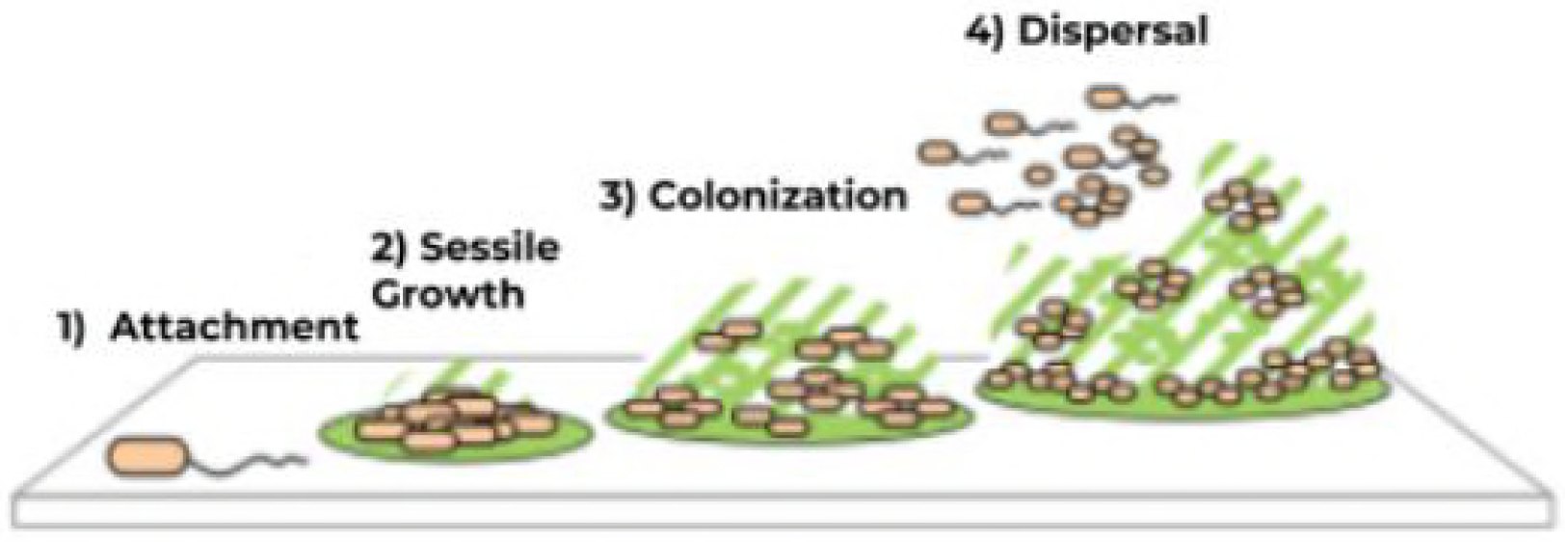
During the attachment stage, the bacteria begin to stick to a surface and start to produce the matrix. After that comes the sessile growth stage, which is when strong adherence and association with other cells develops, and the matrix continues to be constructed, providing protection. Third is the mature colonization stage, when the matrix becomes fully formed, gradients are established, and the bacteria are highly resistant to most antibacterial compounds and conditions. Lastly is dispersal, when bacteria break off from the original biofilm and act as starters for new ones. Adapted from Taylor et al., 2014.

Biofilms are highly resistant to a variety of conditions that would normally kill planktonic cells, most notably antibiotics. There are a variety of techniques that have been developed to attempt to combat this antibiotic resistance (Figure 2). One path being focused on is ways to cause biofilm dispersal, as, upon being restored to their planktonic state, the bacteria are susceptible to antibiotics. This can be done through modification of environmental conditions including pH and oxygen availability, the digestion of the matrix with enzymes like DNase1, or a variety of chemicals like nitric oxide (Taylor et al., 2014).

**Figure 2:**
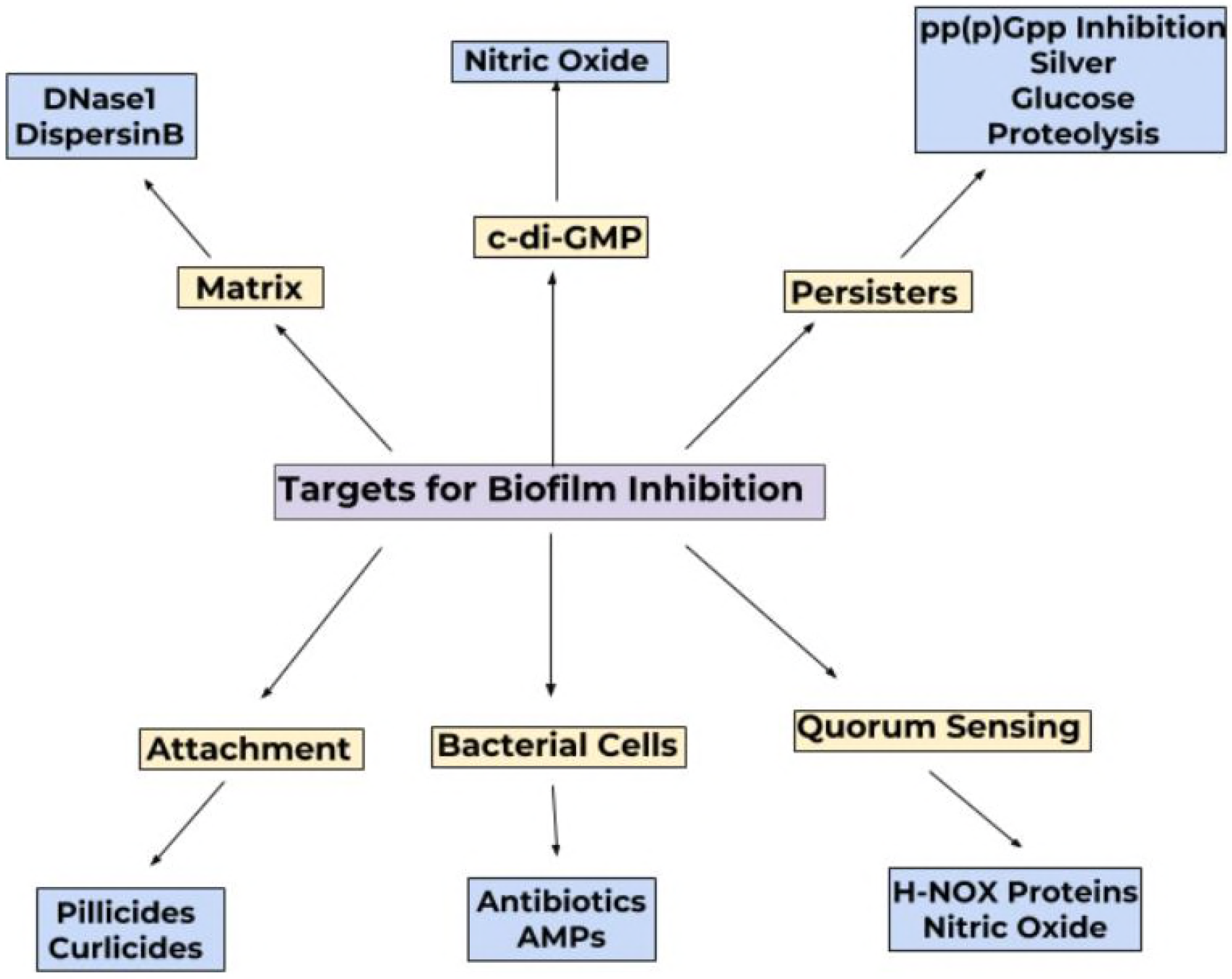
The yellow boxes contain potential features of biofilms that treatments could aim to disrupt. The blue boxes below their respective features are established examples of treatments that target them. AMPs stands for antimicrobial peptides.

One emerging field in biofilm treatment is the use of antimicrobial cationic peptides. These molecules are naturally produced by humans and other animals and have potent, rapid, and broad-spectrum effects against gram-negative and gram-positive bacteria. They have multiple mechanisms of action, making it harder for bacteria to become resistant to them, and can work alone and in combination with antibiotics (Dosler & Karaaslan, 2014). Their amphipathicity combined with their positive charge (as they are cationic), makes it possible for them to interact with membranes, usually by creating pores. However, as natural products they can be unstable, potentially toxic, and costly to produce. Therefore, attempts have been made to design them rationally, in order to optimize their performance and minimize these hindrances (de la Fuente-Núñez et al., 2017). Melittin is one of these antimicrobial peptides; it is found in honey bee venom and works by destabilizing cell membranes, likely through permeabilization, artificial pore formation, disruption and/or lysis (Picoli et al., 2017). It is effective against *Escherichia coli* biofilms, with an MIC of 40-42.5μg/mL and an MBC of 64-128μg/mL (Picoli et al., 2017).

When antimicrobial peptides are used in combination with antibiotics they have been shown to usually have additive or synergistic effects rather than antagonistic, and that using them in combination decreases their minimum biofilm eradication concentration to clinically manageable levels, making them a promising treatment to test in clinical trials (Dosler & Karaaslan, 2014). Melittin is effective against biofilms, but only in large and expensive concentrations that are not feasible for practical use (Picoli et al., 2017). A way that has been shown to increase the efficacy of antimicrobial peptides against biofilms is the usage of them in combination with other antibiofilm agents, as this can create synergistic effects (Dosler & Karaaslan, 2014). DNase I is one potential option for this, as it can disperse biofilms at doses as low as 1 μg/mL (Tetz et al., 2009). This study investigates the synergistic interactions of melittin and DNase I against *E. coli* biofilms.

## Methods

### Biofilm Formation

A bacterial culture was grown overnight in 100 mL of LB broth in a 250mL flask in a shaking water bath using an *Escherichia coli* K12 strain from Carolina Biological. The next day, this culture was diluted in M9 minimal salts supplemented with .2% glucose and 1mM MgSO_4_ using a 1:100 ratio of overnight to M9 by adding 10μL of overnight to 1 mL of M9 in each well of the first five columns of six flat-bottomed 24-well plates. The overnight was regularly vortexed to prevent disparities in *E. coli* concentration. These plates grew at 37°C for three days.

### Treatment Preparation

The melittin (Sigma-Aldrich) was reconstituted and diluted, using sterile deionized H2O for both processes, to a concentration of 84μg/mL, a value roughly double its minimum inhibitory concentration, and the DNase I treatment was reconstituted using water and diluted using PBS to a concentration of 5μg/mL, based upon data regarding its action against *E. coli* biofilms in various doses over time (Picoli et al., 2017; Tetz et al., 2009).

### Biofilm Eradication and Planktonic Bacteria Quantification Assays

The treatments used were 50μl of water (the solvent the melittin is dissolved in) and 50μl of PBS (the solvent the DNase I is dissolved in); 50μl of the melittin; 50μl of the DNase I; 50μl of the DNase I closely followed by 50μl of the melittin; and with 50μl of the DNase I followed by 50μl of the melittin 18 hours later. Four wells were treated with each and then the plates were placed in an incubator at 37°C. In each trial six plates were set up. Data were collected from the first group of three plates after 24 hours had passed since the first treatment was introduced, and after 48 hours for the second group of three plates. To collect the data on planktonic cells, the culture in each well was pipetted out, diluted 0.5μl of overnight to 50mL of LB media and plated. The CFUs of the plates were counted after they were allowed to grow at 37°C for one day. After pouring off the media, the well plates were washed with tap water twice and allowed to dry. The biofilms were then stained by adding 1mL of 0.1% crystal violet solution and left at room temperature to incubate for 15 minutes. Then, the wells were again washed completely with water and allowed to dry. To determine the optical density of the cells, 3 mL of denatured alcohol was added to the plates to solubilize the crystal violet and left for 15 minutes. Then, 1mL of the solubilized crystal violet from each well was be transferred to a cuvette and absorbance was measured using a spectrophotometer at a wavelength of 570 nm. The solubilized dye from a well that had not been inoculated with bacteria was the blank.

### Resistance Development Assay

In order to quantify the variation in bacterial response to the most effective treatmentthe DNase I treatment was tested on 11 bioreplicates. Bioreplicates were prepared by selecting 16 *E. coli* colonies from a stock plate and growing each individually as overnights in LB broth in a shaking water bath at 37C. Then, the OD_600_ of each culture was measured, and the 11 bioreplicates with the highest ODs were selected to be used for the plating. They were all diluted using LB to the OD of the bioreplicate (out of the eleven) with the lowest one, then 10 μL of each overnight was added to 1 mL of M9 minimal salts supplemented with 0.2% glucose and 1mM MgSO_4_ in two wells of a 24-well plate. The plate was grown at 37°C for 72 hours, then the DNase I was applied to each well in the same amount and concentration as described above as it was the most effective treatment from the biofilm eradication assay was applied to them in the same amount and concentration as described above. After 48 hours, data were collected from these plates using the previously described crystal violet assay.

## Results

### Biofilm Eradication

The efficacy of various treatments against *E. coli* biofilms was tested at 24-hour and 48-hour time points (Table 1). The ANOVA indicates that there was significance in the results. Tukey’s test showed that, for the biofilm eradication assay, a significant difference was detected between two groups, specifically the melittin and the DNase I treatments. The difference between melittin and the control was not significant, nor was the difference between DNase I and the control. These findings were consistent at both 24 and 48-hours (Figures 1a and 1b). There was a trend of an increase in biofilm density between the 24- and 48-hour time periods in all treatment groups, but this was not statistically significant.

**Table 1:** ANOVA table for the biofilm eradication assay

| Source | DF | Sum of Squares | Mean Square | F Ratio | P-Value |
| --- | --- | --- | --- | --- | --- |
| Model | 9 | 0.06132987 | 0.006814 | 6.4558 | .0214 |
| Treatment | 4 | 0.01268450 |  | 3.0042 | .0005 |
| Time | 1 | 0.01373880 |  | 13.0158 | .4069 |
| Time*Treatment | 4 | 0.00425378 |  | 1.0075 |  |
| Error | 110 | 0.1161105 | 0.001056 |  |  |
| C. Total | 119 | 0.17744037 |  |  |  |

**Figures 1a and 1b:**
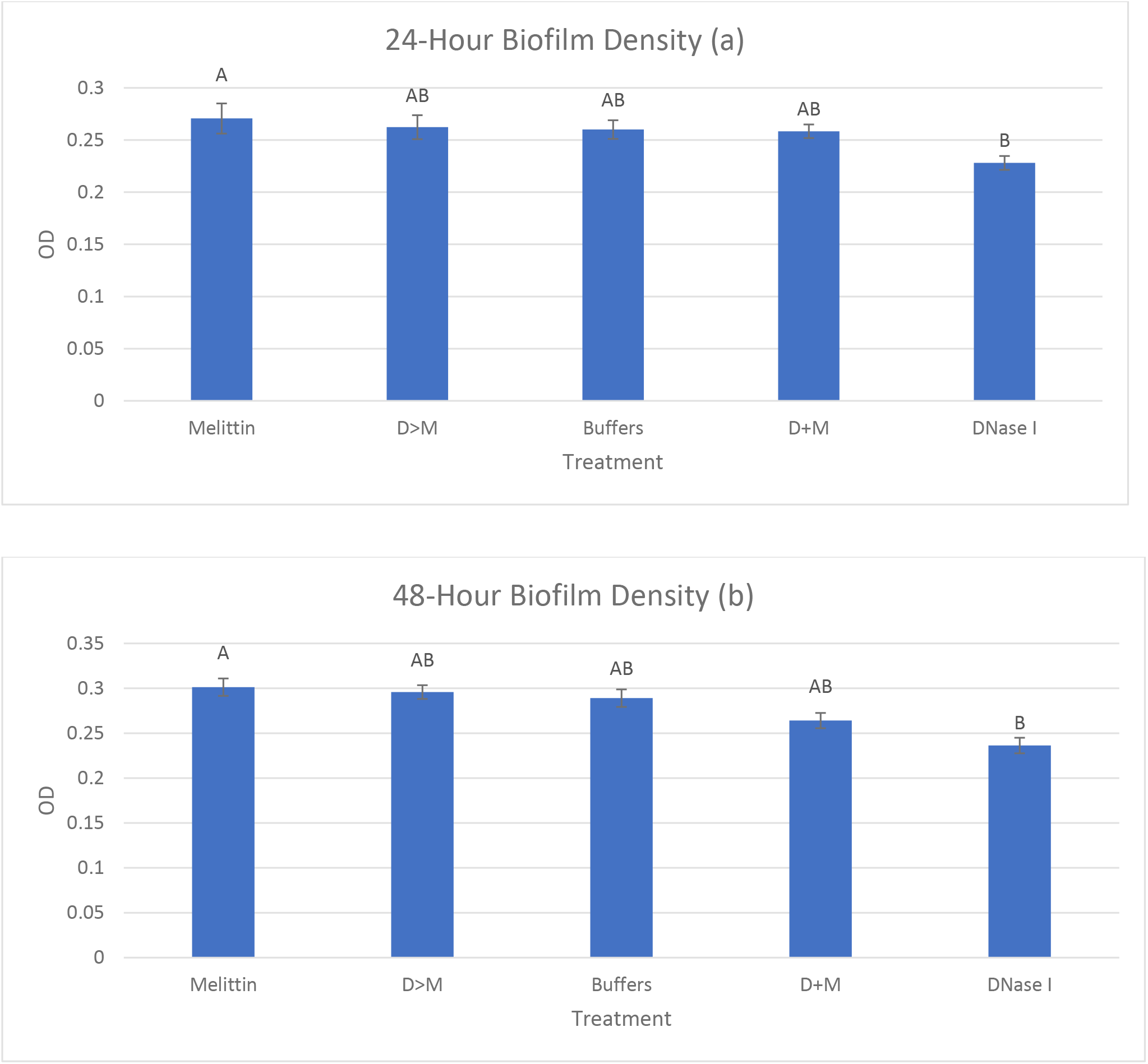
The top (a) is a comparison of biofilm density between treatment groups with letters to indicate statistical significance at 24-hours. The bottom (b) compares biofilm density in the same manner at 48-hours. Both show the biofilms treated with DNase I being significantly less than those treated with melittin (ANOVA p<.001). Letters indicate statistically significant differences. Tukeys, and error bars are one SEM.

### Planktonic Bacteria Quantification Assay

This assay looked at whether any of the treatments caused an increase in planktonic bacteria in the liquid culture of the wells. No significant effect on the biofilm density by the treatments was found in the 24-hour plate, but two standard errors of the mean for the 48-hour CFUs of the DNase I treatment did not overlap with those of the buffer control (Figure 2a). The DNase I treatment was not different from any other treatment in the 48-hour time period though (Figure 2b).

**Figures 2a and 2b:**
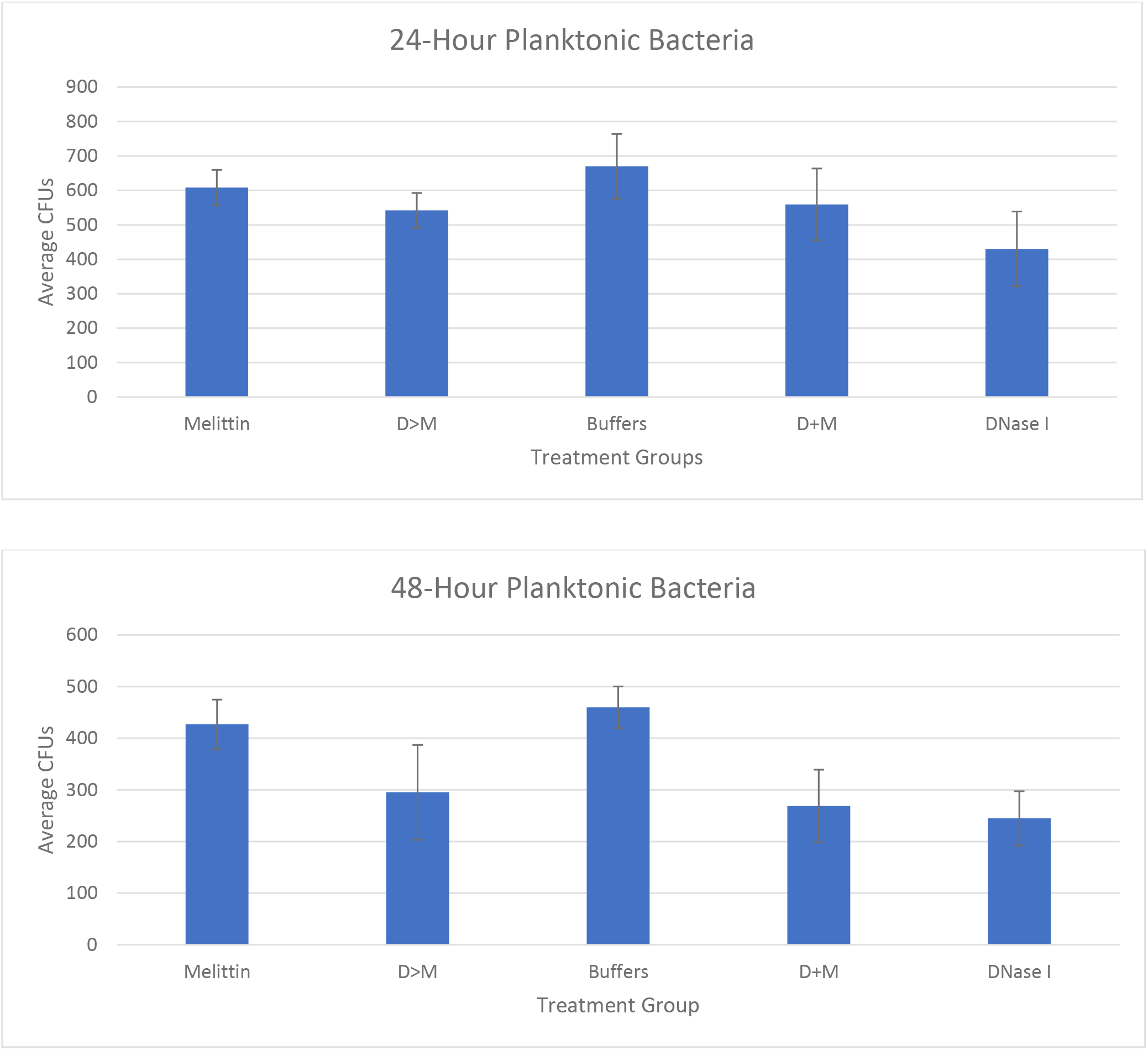
There was (a) no statistically significant difference among any of the treatments in the 24-hour group but (b) two standard errors of the DNase I treatment did not overlap with the buffer treatment at the 48-hour time point. Error bars represent one standard error.

### Resistance Assay

Biological replicates of *E. coli* were treated with DNase I, the most effective treatment from the biofilm eradication assay, to determine if any were unusually resistant to the treatment. There was variance in susceptibility and resistance among biological replicates, more so than within the replicates (Figure 3).

**Figure 3:**
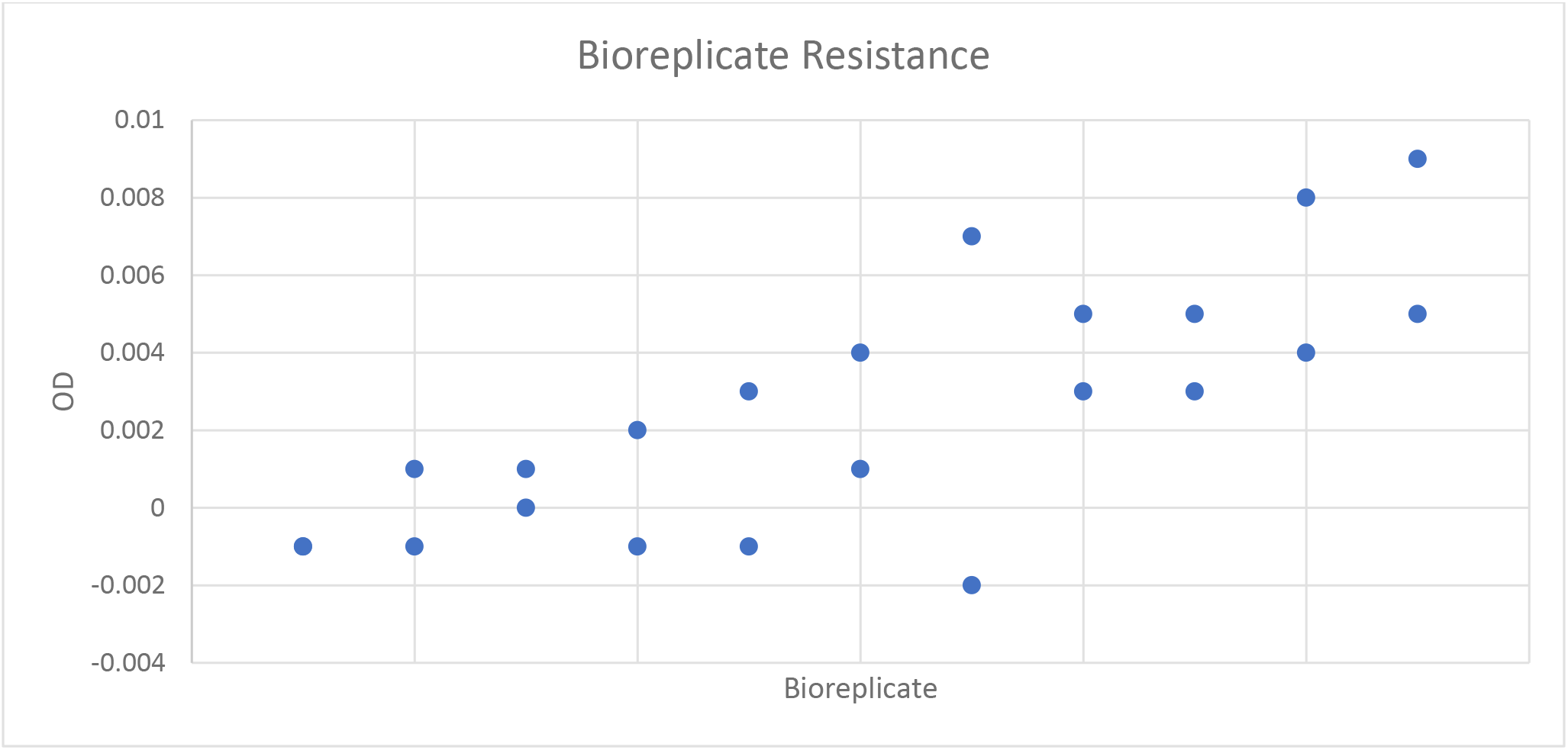
Each point indicates an OD for biofilm quantification after treatment with DNase I, with each biological replicate having two points. This demonstrates greater variance without than within, and an overall low quantity of variance.

## Discussion

The hypothesis behind this experiment was that the DNase I would break down the matrix of the biofilm and allow the melittin greater penetration, and therefore give it greater efficacy. Results showed that melittin was not an effective treatment for *E. coli* biofilms, in contrast to previous literature(Picoli et al., 2017), but that DNase I potentially was. A possible reason for the DNase I being significantly different from the melittin but not from the buffer is that when sub-inhibitory concentrations of treatments were applied to biofilms, they can actually slightly promote biofilm growth (Picoli et al., 2017; Jolivet-Gougeon & Bonnaure-Mallet, 2014). This result indicates that the melittin concentration could have been too low, which could have been caused by mishandling of the protein. The data indicate that DNase I and melittin were not a viable combination treatment, as melittin was generally ineffective, and there was no improvement with that seen in the combination treatments. While the DNase I treatments showed statistically significant difference from the melittin, the combination treatments failed to do so. The use of them together would likely not be a strong treatment for a biofilm infection. One thing that may have caused the DNase I treatment to be not significantly different from the combination treatments, is that the DNase I could have contained peptidases as it was from a bovine source. The length of exposure to the treatments did not influence these results, as the significance was the same at both the 24-hour and 48-hour data collection periods, although the treatments did significantly increase over that time as a whole.

The lack of difference (even slight reduction) in the planktonic bacteria quantification assay was encouraging for the potential use of DNase I for treatment of biofilms though, as it indicates that the concerns over its potential to encourage recolonization may not have a huge impact (Tetz et al., 2009). There was variation among bioreplicates treated with it though, indicating that different strains developing resistance against it is a possibility. DNase I still seems to be a strong potential option for use in clinical settings against biofilms and further research should be done on the use of DNase I in a living subject. Further research should also be done on the use of DNase I with other antimicrobial peptides, to see if it could have synergistic effects with peptides other than melittin and help to expand the field of potential biofilm treatments in that way. In regard to melittin, the effect of DNase I and melittin combination treatment when the positive melittin control is shown to be active against the biofilm as expected should be examined. Melittin should also be tested with other forms of dispersive agents to see if it could function synergistically with those.

## Acknowledgments

I would like to acknowledge Dr. Amy Sheck, NCSSM; Dr. Kim Monahan, NCSSM; Dr. Dan Teague, NCSSM; Tyler Edwards; Kevin Zhang; the Research in Biology Classes of 2018 and 2019; Dr. Daina Zang and the Agile Sciences Lab; and the Glaxo Endowment to NCSSM.

